# Abscisic-acid-dependent regulation of *Arabidopsis thaliana* ammonium transport relies on ABI1 control of CIPK23 and AMT1

**DOI:** 10.1101/2021.02.09.430435

**Authors:** Pascal Ganz, Romano Porras-Murillo, Toyosi Ijato, Jochen Menz, Tatsiana Straub, Nils Stührwohldt, Narges Moradtalab, Uwe Ludewig, Benjamin Neuhäuser

## Abstract

Ammonium uptake at plant roots is regulated at the transcriptional, post-transcriptional and post-translational levels. Phosphorylation by the protein kinase CIPK23 transiently inactivates the ammonium transporters (AMT1s) but the phosphatases activating AMT1s remain unknown. Here, we have identified the PP2C phosphatase ABI1 as an activator of AMTs in *Arabidopsis thaliana*. We show that high external ammonium concentrations elevate the stress phytohormone abscisic acid (ABA) by de-glycosylation. Active ABA is sensed by ABI1-PYL complexes followed by the inactivation of ABI1 activating CIPK23. Under favourable growth conditions, ABI1 reduces AMT1 phosphorylation, both by binding and inactivating CIPK23, and by the direct dephosphorylation of AMT1s. Thus, ABI1 is a positive regulator of ammonium uptake, coupling nutrient acquisition to abiotic stress signalling. Elevated ABA reduces ammonium uptake during stress situations, such as ammonium toxicity, whereas ABI1 reactivates AMT1s under favourable growth conditions.

## Introduction

Uptake of nitrogen from the soil in the form of NO_3_^-^ or NH_4_^+^ is crucial for plant growth and has to be coordinated with the uptake of other ions to maintain cellular ion homeostasis and plasma membrane potential. This complex task is managed on the transcriptional, post-transcriptional and post-translational levels by a variety of stimuli and regulators (Krouk et al. 2010). Whereas ammonium uptake in *Arabidopsis thaliana* is subjected to strong transcriptional regulation by light, sugar and glutamine (Gazzarrini et al. 1999; Rawat et al. 1999), it is additionally regulated and fine-tuned by multiple phosphorylation sites in the AMT1 C-termini (Loqué et al. 2007; Neuhäuser et al. 2007; Straub et al. 2017; Wu et al. 2019). Following the identification of a conserved phosphorylation site in AMT1 C-termini (Nuhse et al. 2004), the phosphorylation of the respective T460 (AMT1;1) and T472 (AMT1;2) residues has been shown to inactivate AMT1 transport activity (Loqué et al. 2007; Neuhäuser et al. 2007). This strong inhibition of uptake has been proposed to be an effective mechanism for avoiding ammonium toxicity (Neuhäuser et al. 2007). In agreement with this hypothesis, the phosphorylation of the AMT1 C-termini shows a fast and strong increase after ammonium resupply and is absent in nitrogen-starved plants (Lanquar et al. 2009; Straub et al. 2017). Adjacent to the conserved threonine residue, the AMT1 C-termini contain several additional non-conserved phosphorylation sites that are differentially phosphorylated upon differences in the nitrogen supply (Engelsberger and Schulze 2012; Menz et al. 2016). In AtAMT1;3, phosphorylation of one of these sites has been shown to fine-tune AtAMT1;3 activity in dependence on nitrate availability (Wu et al. 2019). These non-conserved phosphorylation sites are hypothesized to integrate multiple environmental stimuli with AMT1-mediated ammonium nutrition (Wu et al. 2019).

Whereas the kinases involved in the AMT-specific phosphorylation mechanisms of the non-conserved C-terminal region remain unclear, ammonium-dependent phosphorylation of the conserved C-terminal phosphorylation site has been shown to be mediated by CIPK23 – CBL interacting protein kinase 23 (Straub et al. 2017). Earlier research has highlighted the role of the kinase CIPK23 as an activator of the potassium transporter AKT1 during low K^+^ supply, while it additionally modifies the affinity of the crucial nitrate transporter NPF6;3 (Xu et al. 2006; Ho et al. 2009) and regulates the maintenance of the iron transporter IRT1 at the plasma membrane and its degradation under metal stress (Dubeaux et al. 2018). These processes are carried out via the phosphorylation of the transporters by a CIPK23 – CBL (calcineurin B-like protein) complex interacting with the transporters at the plasma membrane (Lee et al. 2007). This suggests that the kinase is crucial for balancing the cation/anion uptake ratio and for the inorganic nitrogen form being taken up. Additionally, the kinase is also involved in the regulation of ammonium uptake by phosphorylation of AtAMT1;1 and AtAMT1;2, resulting in inactive transporters (Straub et al. 2017). Whether the activity of the AMTs is restored after phosphorylation remains unknown.

The Arabidopsis genome contains approximately 150 protein phosphatases, but not all of them are expressed in roots (Kerk et al. 2008). The *ABI1* gene (*ABA-INSENSITIVE1*) encodes a ubiquitously expressed serine/threonine type 2C protein phosphatase that is a member of clade A in the PP2C phylogenetic tree (Xue et al. 2008). This clade consists of nine members. ABI1 and its homologue ABI2 were early identified as key components of ABA signalling (Leung et al. 1997). Ammonium toxicity has been genetically linked to abscisic acid (ABA) signalling (Li et al. 2012). Mutants of both genes are insensitive to ABA, indicating their role as negative regulators in the pathway (Gosti et al. 1999; Merlot et al. 2001).

Members of the PP2C clade A are regulators of calcium-dependent kinases. The binding of the phosphatase to the kinase prevents (auto-)phosphorylation of the kinase. This interaction is cancelled when the phosphatase comes into contact with an ABA/ABA-receptor complex formed by PYR/PYL proteins (PYRABACTIN RESISTANCE 1/PYRABACTIN RESISTANCE 1–Like) and ABA. Liberation of the kinase finally leads to its activation and target phosphorylation (Park et al. 2009; Umezawa et al. 2009; Yin et al. 2009). The interaction of ABI1 with SNF1, SnRK2.6 (SNF1-related protein kinase 2, OST1) and SnRK2.4 in the absence of ABA is a well-known example leading to kinase inhibition (Yoshida et al. 2006; Lynch et al. 2012; Krzywińska et al. 2016). Furthermore, HAB1 and HAB2 have been shown to interact with OST1 (Vlad et al. 2009). Interestingly, two clade A PP2Cs, namely PP2CA and AIP1 (At1G07430), are both able to regulate the potassium channel AKT1 by modulating CIPK6 and CIPK16 activity (Lee et al. 2007; Lan et al. 2011). Moreover, AIP1 interacts with AKT1 and is therefore proposed directly to inhibit AKT1 activity by dephosphorylation.

In this study, we have identified that the ABI1 phosphatase regulates ammonium transport and have addressed its role in regulating AMT1 activity in an ammonium- and ABA-dependent manner. We hypothesize that ABI1 stimulates AMT1 activity both by direct dephosphorylation and by CIPK23 inactivation and that ammonium toxicity as an abiotic stress is signalled by increased ABA in the roots. Consequently, the binding of a PYL/ABA complex to ABI1 inhibits AMT1 and CIPK23 dephosphorylation by ABI1 resulting in ammonium-dependent phosphorylation and inactivation of AMT1s by CIPK23.

## Material and Methods

### Plant material

Wild-type Col-0, a knock-down mutant of *ABI1* (AT4G26080) *abi1-2* (NASC Nr.: N655606; SALK_072009C), complementation lines generated by agrobacterium-mediated transformation of *abi1-2* with a wild-type *ABI1* coding sequence having the 1000-bp promotor region and previously used AMT / CIPK23 transgenic plants (Straub et al., 2017) were used in all experiments. Additional 73 T-DNA insertion lines close to or in the coding region of phosphatases highly expressed in the root were used in the initial screen for AMT1 activators (Tab. S1).

### Generation of complementation lines

A genomic fragment covering the 1000-bp promotor sequence upstream of the start codon and the open reading frame of *ABI1* were cloned into pTbar vector yielding the transformation vector *pTbar pABI1::ABI1*. Homozygous *abi1-2* plants were transformed with the floral-dip method (Clough and Bent 1998) by using the *Agrobacterium tumefaciens* strain GV3103. Plants were propagated for three generations and two independent homozygous complementation lines were obtained. Selection was performed on ½MS media supplemented with 4 µM L-methionine sulfoximine (MSX). Gene expression of *ABI1* was verified by qRT-qPCR.

### Ammonium uptake assay

A hydroponic system was used to grow the plants for 6 weeks in quarter-strength Hoagland solution at pH 6 with 1 mM NH_4_NO_3_. The controlled environmental conditions were: 8 hours short day (light 200 µmol m^-2^ s^-1^) with a temperature between 20-22°C and a humidity range of 50 to 60 %. After six weeks, plants were transferred to ¼ Hoagland media lacking nitrogen and starved for four days. Afterwards, the plants were transferred to ¼ Hoagland media supplemented with 0.25 mM and 2.5 mM (^15^NH_4_)_2_SO_4_ for 10 minutes. Roots used for ^15^N-measurements were washed twice with 1 mM CaSO_4_, briefly dried with a paper towel, frozen and freeze-dried. Plants for the second time-point were placed in shocking solution containing 10 mM (NH_4_)_2_SO_4_ for five minutes and harvested after uptake in the same way as that described above at 2.5 hours after the beginning of the shock. The dried plant material was ground and 0.5 mg was taken to measure the δ ^15/14^N ratio by using isotope ratio mass spectrometry (Delta plus Advantage, Thermo Fisher Scientific, coupled to an Eurovector elemental analyser EA 3000).

### Gene expression analysis

Samples were taken at each harvesting time-point of the ammonium uptake assay and immediately frozen in liquid nitrogen. Afterwards, they were ground using mortar and pestle and five plants were pooled as one biological sample. Total RNA was extracted with the innuPREP Plant Extraction kit from Analytik Jena and the concentration was measured using a NanoDrop2000 spectrophotometer (Thermo Fisher Scientific Inc., Waltham). Reverse transcription was performed with the QuantiTect Reverse Transcription Kit (Qiagen GmbH, Hilden) according to the manufacturer’s manual. The qRT-PCR was set up with the GreenMasterMix (GENAXXON bioscience, Ulm) and a C1000 Thermo Cycler attached to a CFX 384 real-time system (Bio-Rad Laboratories Ltd., Watford). Each reaction volume of 15 μL contained 10 ng cDNA. Relative expression was calculated using the ΔΔCT method with UBQ10 and SAND as reference genes. The primers used are shown in Table S2.

### Yeast two-hybrid interaction assay

The coding sequences of *ABI1, AMT1;1, AMT1,2, CIPK23* and all *PRY/PYL* genes were amplified from cDNA and cloned into pCR™Blunt II-TOPO® vector (Thermo Fisher Scientific, Waltham) followed by amplification with restriction-site-containing primers and subcloning into the corresponding vectors as described below. The open reading frame of *ABI1* was cloned into pPR3-N, whereas *AMT1;1, AMT1;2*, their C-terminal mutants and *CIPK23* were ligated into pBT3-C, both vectors being part of the DUALmembrane Starter Kit (Dualsystems Biotech AG, Schlieren). The *PRY/PYL* genes were also cloned into pPR3-N. Combinations of both vectors were transformed into yeast strain NMY51 by the lithium acetate method (Gietz and Schiestl 2007). Positive transformants were selected on solid SD-Trp-Leu medium. Colonies for the assay were picked and grown overnight in liquid SD-Trp-Leu at 28°C. Afterwards, they were washed three times with water and adjusted to an OD_595_ = 2. Yeast suspension in a volume of 10 µL was spotted onto SD plates without -Trp-Leu-Ade, -Trp-Leu-His or -Trp-Leu-Ade-His; for ABI1/PYL interaction, they were further supplemented with 10 µM ABA. Yeast was grown for 5 days at 28°C and were subsequently covered with an X-Gal overlay. The mutants *AMT1;1ΔS488-V501* and *AMT1;1ΔT497-V501* were generated by site-directed mutagenesis PCR resulting in a deleted C-terminus followed by a stop codon.

### Split-YFP interaction in planta

The *in planta* interaction between ABI1 and AMT1;1/AMT1;2/CIPK23 was tested as previously described (Straub et al. 2017). With regard to the design of the fusion construct, the YFP molecule was split between amino acids 153 and 154 and these parts were inserted into the pTkan^+^ and pTbar vectors to give the pTkan^+^YN 153stop and pTbar YC stop vectors. The vectors containing the AMT1s and CIPK23 were available in our laboratory. The respective promotor:gene fusion for ABI1 was cloned into the two vectors by using appropriate single cutting enzymes in the MCS. The various plasmid combinations were transformed into *Arabidopsis thaliana* Col-0 plants as described above. Transgenic plants were analysed by using a LSM700_ZEN_2010 microscope (Zeiss, Germany).

### Protein extraction and immunoblotting

Total protein was extracted from frozen root tissue by using cold extraction buffer (100 mM NaCl, 50 mM Tris/HCl (pH = 7.5), 0.5 % (v/v) Triton X-100, 10 mM β-mercaptoethanol) supplemented with PhosSTOP (Roche, Basel) and Complete Protease Inhibitor Cocktail (Roche, Basel) according to the manufacturer’s manual. Samples were then centrifuged and the protein concentration in the collected supernatants was measured using the Bradford assay. Proteins in amounts of 10 µg were denatured in Laemmli sample buffer (62.5 mM TRIS pH 6.8, 2 % (w/v) SDS, 10 % (v/v) glycerol, 5 % (v/v) β-mercaptoethanol, 0.001 % (w/v) bromphenol-blue) for 10 minutes at 50 °C. Subsequently, samples were loaded onto a 12 % acrylamide SDS gel and separated by electrophoresis. Transfer to a nitrocellulose membrane was performed by semi-dry blotting with standard Bjerrum Schafer-Nielsen buffer (Bio-Rad Laboratories GmbH, Feldkirchen). Membranes were blocked using TBS-T containing 1 % (w/v) casein hydrolysate for 1 hour followed by overnight incubation in blocking solution and primary antibody (IgG, rabbit, dilution 1:1000; BioGenes, Berlin). The custom-made antibody detected the phosphorylated conserved threonine in AMT1 (CG-NleD-Nle-pT-RHGGFA-amide, Fig. 3b). After 3 washing steps, the membrane was incubated for 1 hour in TBS-T supplemented with the secondary antibody (polyclonal IgG, goat, conjugated to horseradish peroxidase, dilution 1:5000; Roth, Karlsruhe). The membrane was subsequently washed with TBS and treated with ECL SuperSignal West Dura solution (Thermo Fisher Scientific, Waltham). Signals were detected on an Odyssey Fc imager (Li-COR Biotechnology; Bad Homburg). The intensity of the bands was measured using ImageJ software. For a loading control, the membrane was incubated for 2 min in 0.1 % (w/v) Ponceau S in 5 % (v/v) acetic acid. Afterwards, the membrane was washed twice in distilled H_2_O.

### Hypocotyl elongation assay

Sterilized Arabidopsis seeds were distributed on square petri dishes containing MQ water supplemented with 20 g/L agar and 40 mM methylammonium (MeA) or 0 mM MeA. The plates were incubated at 4 °C for 2 days. Germination was induced by incubation in a Percival AR-66L (150 μmol m^-2^s^-1^, Philips F17T8/TL841 17W) for 1 day, followed by 5 days incubation in the dark. Hypocotyl length was measured using ImageJ software and the growth reduction of the *abi1-2* mutant and the complementation lines was normalized to control conditions and/or Col-0’s hypocotyl length. The experiment was repeated three times; approximately 30 hypocotyls were measured per treatment, line and repetition.

### Abscisic acid measurements

Frozen Arabidopsis roots were ground to a fine powder with liquid nitrogen and 1 g plant material was treated with 80 % methanol. Supernatants were collected after centrifugation and cleaned by membrane filtration (Chromafil® O-20/15 MS). Samples were then analysed by UHPLC-MS on a Velos LTQ System (Thermo Fisher Scientific, Waltham) as described in more detail by Moradtalab and co-workers (Moradtalab et al. 2018).

### Statistics

At least three repetitions were conducted in all experiments. The numbers of the replicates in one experiment are given in the figure legends. Statistical significance was tested by an ANOVA followed by Tukey’s post-hoc test. Significant differences are indicated by capital or small letters (p < 0.5). When the ANOVA was followed by a pairwise comparison, significant differences are indicated by *** for p < 0.001.

## Results

### Knockdown mutant abi1-2 shows reduced methylammonium susceptibility

To identify possible AMT1 activators, we selected phosphatases showing at least moderate expressed in roots. With regard to phosphatase families with several candidates, we focused on those that were most strongly expressed (Tab. S1). In an initial screen of 73 *Arabidopsis thaliana* T-DNA insertion lines, we tested these root-expressed phosphatase candidates for reduced sensitivity to toxic ammonium and methylammonium (MeA) conditions. Because of the high variability of the root growth, we initially confirmed that the candidates were also expressed in the hypocotyl; we proceeded with a hypocotyl elongation assay and identified the *abi1-2* (from here on *abi1*) knock-down mutant (Wu et al. 2015) and thereby ABI1 (At4G26080) as a potential regulator of AMT1 activity. We first confirmed that the T-DNA insertion in *ABI1* (At4G26080; SALK_072009C) was homozygous and resulted in an approximate 90 % knockdown of *ABI1* expression (Fig. S1a/b). Second, we reproduced the initial screening test (hypocotyl elongation) for Col-0 and *abi1* with an increased number of replicates (n > 200). This experiment confirmed the reduced ammonium and methylammonium susceptibility of the *abi1* line (Fig. S2b). T-DNA insertion lines for three other clade A members were also part of our screen (*HAB1*: At1G72770; *HAB2*: At1G17550 and *PP2CA*: At3G11410); however, these did not confer reduced NH_4_^+^ susceptibility, suggesting specificity of ABI1 in this regulation (Fig. S2a).

To establish the specificity of the effect in the *abi1* line, we created two independent complementation lines; these slightly overexpressed *ABI1* and from here on are referred to as OX1 and OX2. All lines were again tested under toxic MeA conditions in the hypocotyl elongation assay (Fig. 1a/b). The mutant *abi1* line again showed longer hypocotyls under toxic methylammonium conditions compared with those of Col-0 plants, whereas both complementation (OX) lines suffered from slightly increased susceptibility (Fig. 1a/b).

**Figure 1.**
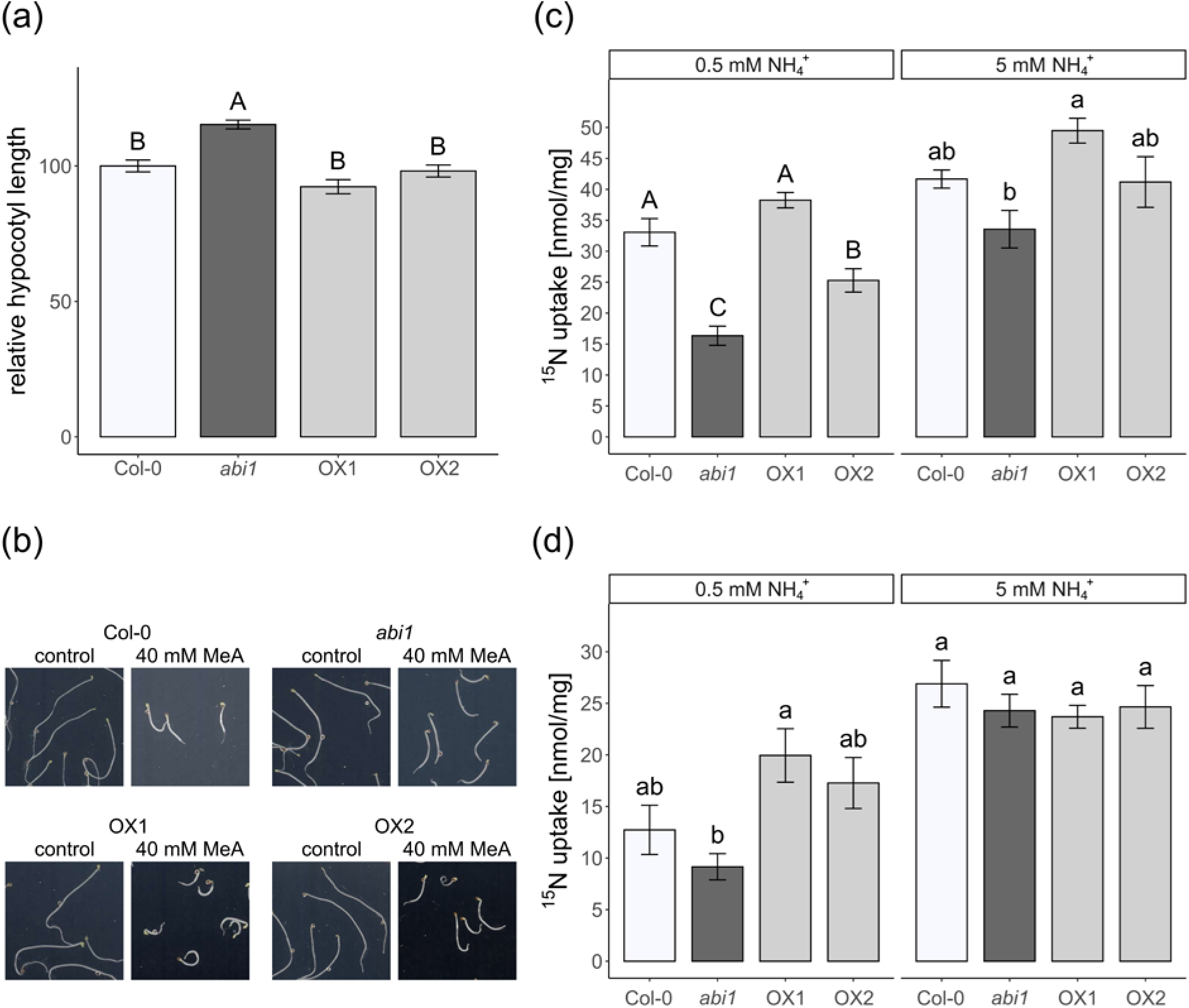
Lower ammonium sensitivity and uptake in *abi1* plants. **(a)** Normalized hypocotyl length [%] of etiolated Col-0, *abi1* and complementation line seedlings grown on media containing 40 mM methylammonium. Relative length ±SEM of three replicates (n ≥ 30). **(b)** Representative images of etiolated seedlings grown under control conditions and in 40 mM methylammonium. Height and width of the images correspond to 20 mm. **(c)** Short term ammonium (^15^NH_4_^+^) uptake (10 min) directly after nitrogen starvation phase. Data are shown as mean ammonium uptake ±SEM (n ≥ 4). **(d)** Short term ammonium (^15^NH_4_^+^) uptake (10 min) at 2.5 h after 5-min ammonium shock to induce AMT1 inactivation. Data are shown as mean ammonium uptake ±SEM (n ≥ 4).

### Nitrogen concentration is slightly reduced in the abi1 shoots

To address the ammonium-sensitive phenotype of the *abi1* line further, we grew wild-type and *abi1* plants under standard hydroponic conditions supplemented with 2 mM ammonium nitrate for 6 weeks (Fig. S3e). Root and shoot fresh weights were similar for WT and *abi1* plants, whereas root dry weight was slightly higher in *abi1* plants (Fig. S3a/b). The shoot nitrogen content was similar but the nitrogen concentration in the *abi1* shoots was reduced (Fig. S3c/d), possibly indicating a reduced ammonium uptake or root to shoot translocation even under standard growth conditions.

### Ammonium uptake in the abi1 line is impaired

To detect the reason for the increased tolerance of *abi1* to toxic ammonium and methylammonium, physiological ammonium absorption in plants roots was examined by short-term uptake. Plants were grown hydroponically for 6 weeks and subsequently placed in nitrogen-deficient solution for 4 days. This period of nitrogen starvation has previously been shown to increase the amount of AMT transcripts and, at the same time, to decrease the phosphorylation of the transporters (Yuan et al. 2007; Lanquar et al. 2009; Straub et al. 2017); a combination of the two effects results in a high ammonium uptake capacity. Starved plants were placed in 0.5 and 5 mM ^15^NH_4_^+^ uptake solution for 10 minutes and the uptake was quantified by isotope ratio mass spectrometry. To induce ammonium-dependent phosphorylation of AMT1s by CIPK23, a second timepoint for nitrogen uptake analysis was set at 2.5 hours after an ammonium shock (20 mM NH_4_^+^) for 5 min, followed by N-starvation conditions.

After starvation, most AMTs in the wild-type and in the complementation lines were expected to be fully de-phosphorylated and active, whereas in the *abi1* line, we expected decreased uptake attributable to existing AMT1 phosphorylation. Accordingly, a lack of dephosphorylation might account for the more than 40 % lower ammonium uptake after starvation in the *abi1* line compared with the wild type (Fig. 1c/d). Furthermore, this would imply hyperactivity of the CIPK23 kinase leading to AMT1 phosphorylation and inactivation. By subjecting the plants to an ammonium shock, we induced AMT inactivation by CIPK23-dependent phosphorylation, followed by N starvation for reactivation after some time. We then quantified ammonium acquisition after 2.5 additional hours of starvation following the shock. We found a tendency towards a reduced uptake in the *abi1* plants after the ammonium shock but this difference was not statistically significant in repeated experiments (Fig. 1c/d). Both complementation lines, however, showed higher ammonium uptake than the mutant line under both conditions. In agreement with the potential role of ABI1 in the regulation of the high affinity ammonium transport, we did not detect any statistically significant differences in uptake in the low affinity ammonium range. The *abi1* line also showed a slight but non-significant decrease in uptake after starvation in the 5 mM samples (Fig. 1d), in agreement with the finding that uptake at this high concentration is the sum of high and low affinity ammonium transporters.

### ABI1 interacts with AMT1;1, AMT1;2 and CIPK23

The reduced ammonium uptake in *abi1* after nitrogen starvation might be related to higher AMT1 phosphorylation under these conditions. This might be explained by a deregulated and hyperactive CIPK23 kinase or by the inability to dephosphorylate AMT1. In both cases, ABI1 needs to interact with CIPK23 or with the AMT1s. We therefore checked whether ABI1 interacted with AMT1s and/or CIPK23.

The *in planta* bimolecular fluorescence complementation assay (BiFC) and the split ubiquitin yeast system were used to address these protein-protein interactions. BiFC fusion proteins containing either the N- or C-terminal part of YFP were expressed using their endogenous promoters. Wild-type control plants did not exhibit any fluorescence, whereas combinations of *ABI1-NY* with *CIPK23-CY* and with *AMT1;1-CY* and *AMT1;2-CY* all exhibited plasma-membrane-localized YFP fluorescence (Fig. 2a). Interaction of ABI1 with CIPK23 and the two AMT1s was also confirmed by using a yeast split ubiquitin assay. Yeast expressing all protein combinations showed growth on control media, whereas the ABI1/CIPK23 and ABI1/AMT1 combinations also promoted growth on selective media, confirming the protein interactions (Fig. 2c).

**Figure 2.**
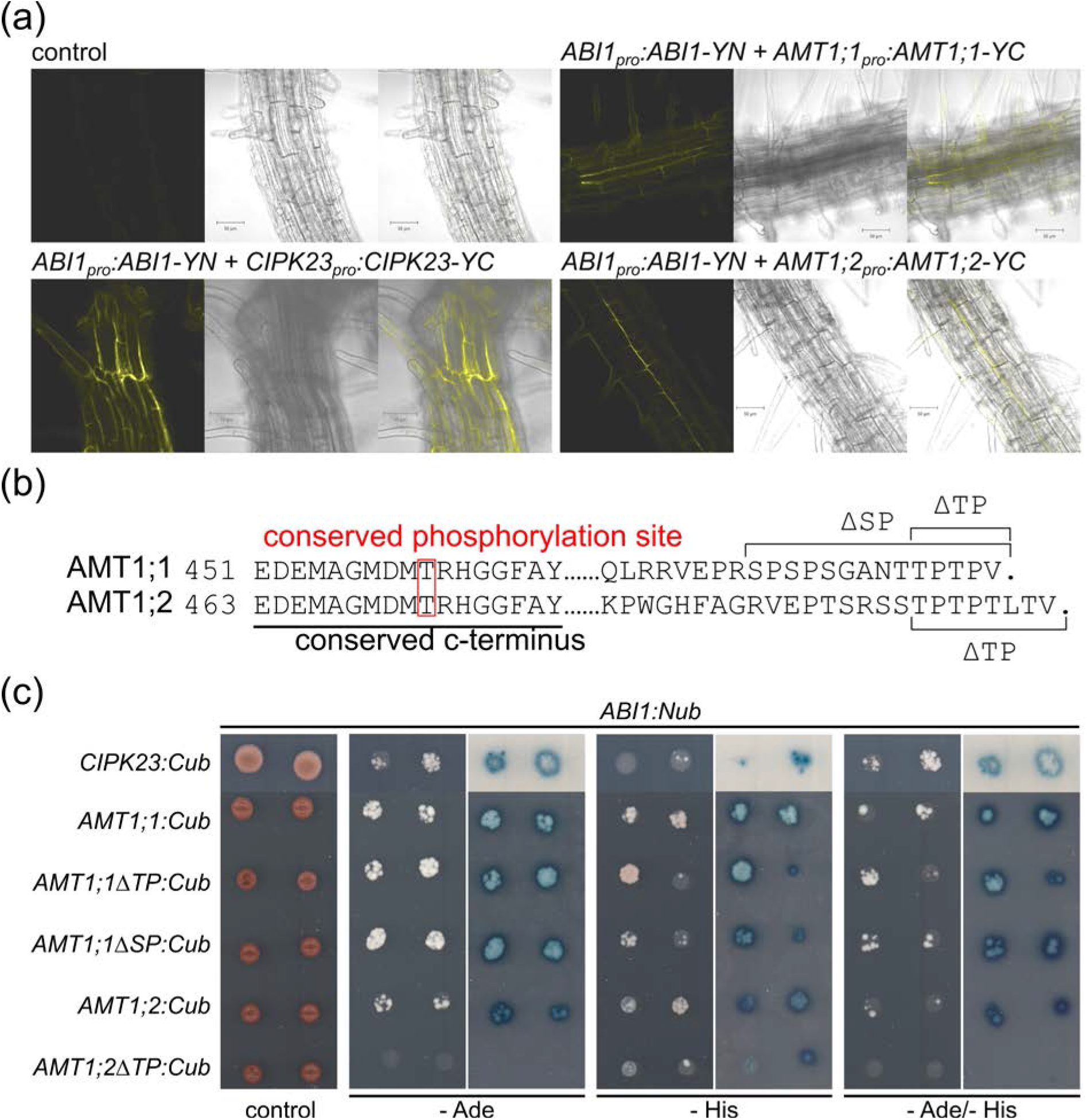
Interaction of ABI1 with CIPK23 and AMT1s. **(a)** Representative confocal microscopy pictures of plants stably co-expressing ABI1, AMTs and CIPK23 fusions carrying split-YFP parts. *Upper left*, Col-0 control; *lower left*, ABI1/CIPK23 interaction; *upper right*, ABI1/AMT1;1 interaction; *lower right*, ABI1/AMT1;2 interaction. From left to right, BiFC-YFP fluorescence signal, brightfield image, overlay. **(b)** Sequence alignment of the AMT1;1 and AMT1;2 C-termini. The conserved phosphorylated threonine residue is highlighted in red. Black line indicates the end part of the conserved C-terminal region. Brackets indicate parts removed in the C-terminal truncation mutants. **(c)** Protein interaction of CIPK23, native AMT1s and C-terminal AMT1 truncation mutants with ABI1. ABI1 was fused to the N-terminal part of ubiquitin, whereas the interaction partners are fused to the C-terminal part of ubiquitin. Growth on media without adenine and/or histidine and blue staining by X-Gal overlay indicate protein interaction. Figure shows one representative image from 3 repetitions.

### ABI1 interacts with AMT1;1 at the conserved C-termini

Several serine and threonine residues in phosphorylation motifs have been found to be differentially phosphorylated in the AMT1 C-termini (Engelsberger and Schulze 2012; Wu et al. 2019). Although only a single threonine residue is phosphorylated in the conserved part of the C-termini, stretches with phosphorylation motifs occur in the non-conserved AMT1;1 and AMT1;2 protein tails (Fig. 2b). The first motif in AMT1;1 (TPTP) starts at position T497, and the second (SPSPS) starts at S488. In AMT1;2, the TPTP motif starts at residue T507. To address which C-terminal segments of the AMTs are of importance for ABI1-binding and might therefore also contain the de-phosphorylation sites targeted by ABI1, we stepwise deleted the respective motifs (Fig. 2b).

We then conducted a split-ubiquitin yeast assay to address the effect of the mutations on the ABI1/AMT1 interaction. As indicated above, protein interaction occurred between ABI1 and both native AMTs in the assay (Fig. 2c), in agreement with the observations made during the BiFC experiment. Interestingly, the C-terminal deletions in AMT1;1 did not disrupt protein interactions. Interaction of native AMT1;2 with ABI1, however, was much weaker compared with AMT1;1 in this assay. Interestingly, the ΔTP deletion of the C-terminus of AMT1;2 resulted in a loss of interaction in yeast (Fig. 2c).

### ABI1 specifically affects the phosphorylation state at the conserved AMT1 C-terminus

Since we could not detect any unspecific effects on *AMT1* expression or AMT1 localization, we concluded that AMT1 phosphorylation should directly be affected in the *abi1* line. To assess the phosphorylation status of the AMT1s at both uptake timepoints, we extracted total root protein followed by phosphorylation-sensitive immunodetection of the conserved AMT1 C-terminus. A custom-made phosphorylation-sensitive antibody targeting the threonine in the conserved C-terminal peptide GMDMT(p)RHGGFA of AMT1s was used to check for phosphorylation differences among the lines.

After four days of nitrogen starvation, we were still able to detect phosphorylated AMTs in all lines after 10 minutes of exposure in an Odyssey Fc chamber (Fig. 3a). The strongest phosphorylation was observable in line *abi1* followed by wild-type Col-0. Both complementation lines showed less AMT1 phosphorylation than Col-0, with line OX1 having the lowest phosphorylation (Fig. S4a).

**Figure 3.**
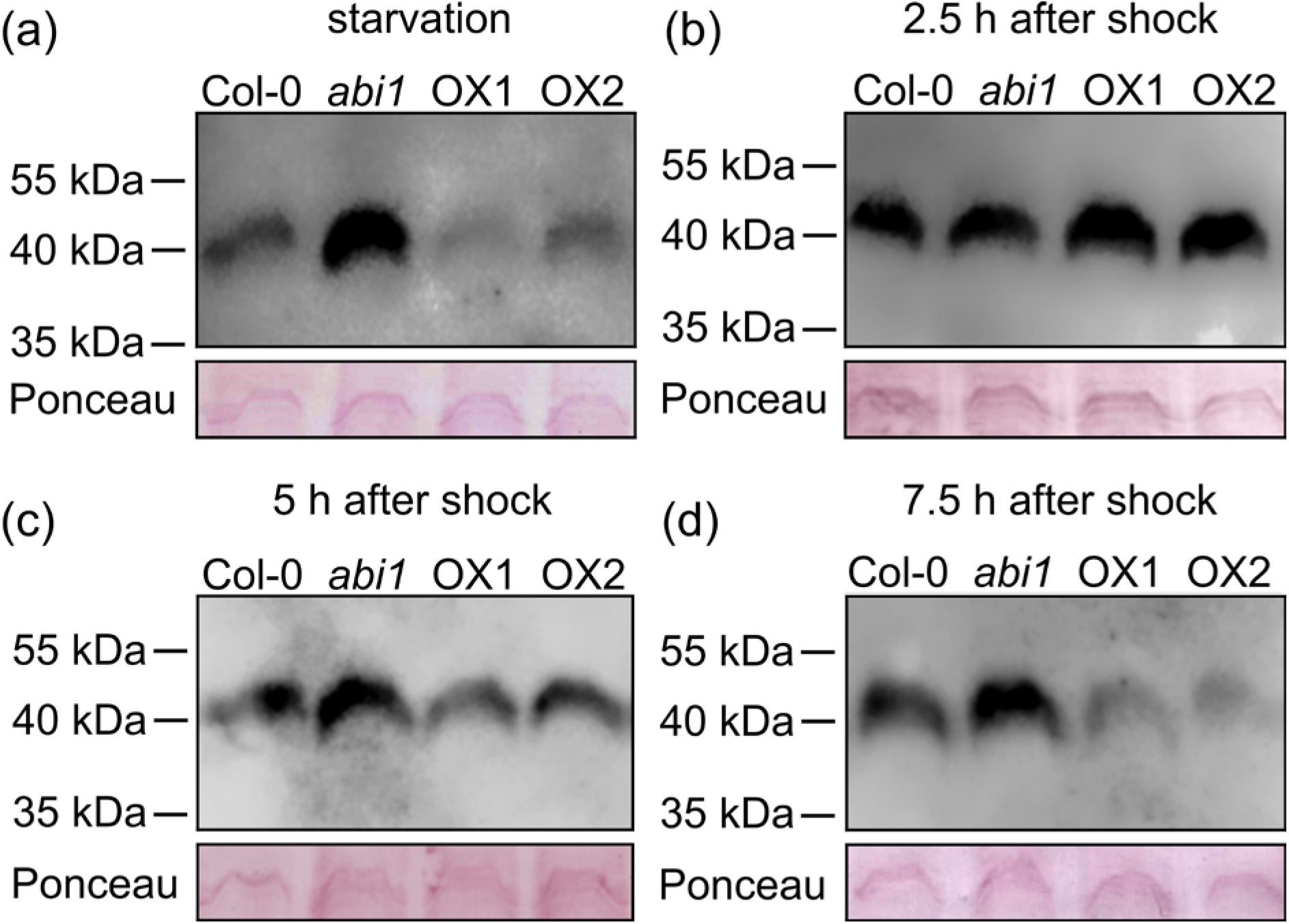
ABI1 affects the phosphorylation status of the conserved AMT1 C-termini. Protein gel blot analysis of total root protein extract from Col-0, *abi1* and complementation lines by using a phosphorylation-specific antibody detecting phosphorylation of the conserved threonine in the AMT1 C-termini after nitrogen starvation for four days **(a)**, after a nitrogen shock followed by 2.5 h **(b)**, 5 h **(c)** or 7.5 h **(d)** of nitrogen starvation. Upper part shows detection of phosphorylated AMT1 C-termini and lower part shows loading controls stained with Ponceau red.

The AMT1 phosphorylation of plants 2.5 hours after the 5-min ammonium shock was completely different. In agreement with previous analyses, the AMT1s were heavily phosphorylated by the shock and this remained in all lines after 2.5 hours of additional starvation period (Fig. 3b; Fig S4b).

At later time points at 5 and 7.5 hours after the shock, however, AMT1s were progressively de-phosphorylated, although this depended on the genetic background. At the 5-hour timepoint, AMT1 dephosphorylation was slightly more pronounced in the complementation lines than in the wild type, whereas AMT1s on the *abi1* background were still heavily phosphorylated (Fig. 3c; Fig. S4c). At 7.5 hours after the shock, we observed the same phosphorylation pattern as that at the beginning of the experiment (Fig. 3d; Fig. S4d), with AMT1s in *abi1* being strongly phosphorylated.

Altered gene expression of the ammonium transporters or their regulatory genes *CBL1* and *CIPK23* might also account for uptake and phosphorylation differences between the lines. However, qRT-PCR of these genes confirmed that no major changes in gene expression occurred under these conditions. This was confirmed for all relevant genes in all plant lines at all investigated time points (Fig. S5). We further ensured that AMT1 localization and protein stability were not affected in the *abi1* plants by checking the fluorescence from AMT1;1-GFP and AMT1;2-GFP fusion constructs expressed under their endogenous AMT1 promoters on the *abi1* and Col-0 backgrounds. On both backgrounds, we observed similar AMT-specific plasma membrane localization and GFP signal strength (Fig. S6).

### Ammonium toxicity increased ABA levels by de-glycosylation

Since ABI1 activity is regulated by ABI1/ABA/PYL complex formation, we analysed whether ammonium toxicity, like other abiotic stresses, is also signalled by ABA. The ammonium shock performed in the uptake experiment led to a similar increase of ABA in the roots. On comparing the ABA concentration in the roots under nitrogen starvation and at the 2.5-hour timepoint, a 9-fold increase after the shock was discovered (Fig. 4a). We then tested whether an external ABA supply could directly affect the short-term ammonium uptake of wild-type plants. Col-0 plants were grown as described above and, after a four-day starvation period, we exposed the plants to 100 µM ABA for one hour, after which we quantified the ^15^NH_4_^+^ uptake at 0.5 mM for 10 min. Equivalent to the *abi1* knockout, the ABA shock strongly reduced the ammonium uptake of the plants (Fig. 4b). Since this ABA increase appeared rapidly, namely only 2.5 hours after the ammonium shock, we next addressed whether it might be the consequence of ABA activation by de-glycosylation. Therefore, we monitored the expression of the *BGLU* genes involved in ABA de-glycosylation in the endoplasmic reticulum and in the vacuole (Ma et al. 2018). After shocking the plants for one hour with 10 mM NH_4_^+^, we observed an increased expression of *BGLU10* mediating ABA de-glycosylation in the vacuole and of its closest homolog *BGLU11. BLGU33 (BG2)* is also important for vacuolar ABA activation but did not increase in expression. *BGLU18 (BG1)*, which is active in the ER, showed an ammonium-dependent increase in expression (Fig. 4c). We finally tested the interaction of ABI1 and the PYR/PYL ABA-receptor proteins in yeast and identified an interaction between ABI1 and almost all PYL proteins. This interaction tended to increase when we added ABA (Fig. S7).

**Figure 4.**
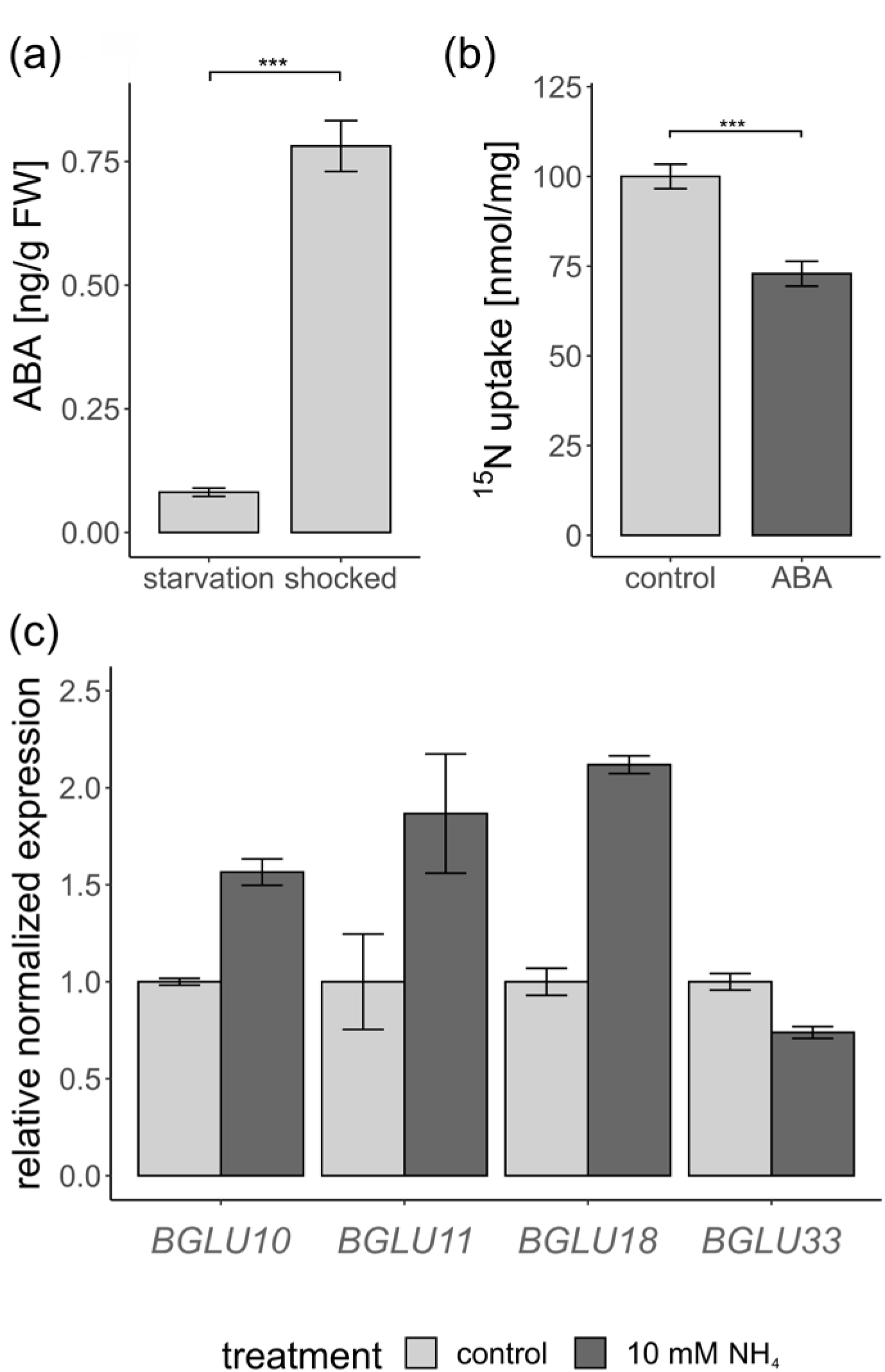
Abscisic acid (ABA) signals are involved in the response to toxic ammonium concentrations. **(a)** ABA concentration ±SEM (n = 8) in starved and shocked plant roots. **(b)** Short-term ammonium (^15^NH_4_^+^) uptake (10 min, 0.5 mM ^15^NH_4_^+^) in Col-0 plants after preincubation in 100 µM ABA. Data are shown as mean ammonium uptake ±SEM (n = 19). **(c)** Normalized expression of *BGLU* genes in starved and shocked plant roots (n = 3).

## Discussion

The activity of many nutrient transporters in plants is balanced by a complex network of phosphorylation and dephosphorylation (Saito and Uozumi 2020). Since ammonium toxicity is detrimental for most plants, a phospho-regulation network acting on plant ammonium transporters provides fast and efficient means to avoid this toxicity. Furthermore, the nutrition of plants has to be adjusted to their physiological status and to any abiotic stresses for acclimatization to environmental conditions as mediated via phytohormone signalling, including the stress hormone ABA (Li et al. 2014; Liu and Von Wirén 2017; Marino and Moran 2019).

Phosphorylation at a conserved C-terminal threonine inactivates plant AMT1 transporters (Loqué et al. 2007; Neuhäuser et al. 2007). However, adjacent to this conserved C-terminal phosphorylation site, AMT1 C-termini harbour several additional non-conserved phosphorylation sites, potentially integrating diverse local and systemic signals with ammonium nutrition (Menz et al. 2016; Wu et al. 2019).

The CBL-binding protein kinase CIPK23 has been shown to regulate the uptake of ammonium, potassium and nitrate by its potential to phosphorylate several important transporters for these nutrients (Xu et al. 2006; Ho et al. 2009; Straub et al. 2017). The CIPK23-dependent phosphorylation of AMT1 transporters at the conserved C-terminal threonine results in a total inhibition of the trimeric AMTs (Loqué et al. 2007; Neuhäuser et al. 2007). Reactivation of the transporters occurs only after complete dephosphorylation (Neuhäuser et al. 2007). This dephosphorylation requires a phosphatase counterpart to CIPK23. Here, we have identified ABI1 as regulating AMT1 activity by direct dephosphorylation, in addition to CIPK23 binding and subsequent inactivation.

ABI1 belongs to the A-subfamily of PP2C phosphatases and is a well-known negative regulator of the ABA response (Rodriguez 1998; Schweighofer et al. 2004). Abiotic stress increases the intracellular ABA concentration, which causes the formation of PYL/ABA/ABI1 complexes and finally leads to ABI1 degradation (Park et al. 2009; Umezawa et al. 2009; Yin et al. 2009). Our data suggest that this complex formation is directly integrated into AMT1 regulation, since plants respond to an ammonium shock by an ABA increase – a mechanism previously established as being a signal of other abiotic stresses (Ma et al. 2018; Chen et al. 2020). A short exposure to high external ammonium concentrations is sufficient to provoke a fast and strong ABA increase in the plant roots (Fig. 4a). Identification of the ammonium over-sensitive mutant AMOS1/EGY1 had previously predicted a connection between ammonium toxicity and ABA signalling (Li et al. 2012). Ammonium has also been implicated in ABA accumulation and root-to-shoot ABA translocation (Peuke et al. 1998; Li et al. 2012; Ding et al. 2016; Liu and Von Wirén 2017). By mimicking this internal ABA increase, the external supply of ABA reduces ammonium uptake to a similar extent as the knockdown of *ABI1* (Fig. 4b and Fig. 1c). Therefore, the external supply of ABA might lead to the direct inactivation or degradation of ABI1. The internal ABA increase occurs rapidly at 2.5 hours after the ammonium shock, which lasts for only 5 minutes. This is coherent with the expression increase of *BGLU* genes involved in ABA de-glycosylation (Fig. 4c). De-conjugation of glucose from the ABA-glycosyl-ester not only increases its bioactivity, but also changes its subcellular localization from the ER and vacuole to the cytosol (Dietz et al. 2000; Kim et al. 2013). BGLU10 and BGLU11 are involved in activating ABA from the vacuole, whereas their close homologue BGLU18 mediates ABA de-glycosylation in the ER (Xu et al. 2012; Ma et al. 2018). Their gene expression increases after the ammonium shock.

As shown previously, we have confirmed that external ABA application increases the interaction of ABI1 with almost all PYL proteins (Fig. S7) (Ma et al. 2009; Park et al. 2009; Yin et al. 2009; Nishimura et al. 2010). Therefore, we propose that an excessive external supply of ammonium will trigger ABA deglycosylation in the vacuole and ER, which in turn leads to PYL/ABA/ABI1 complex formation and AMT1 inactivation by phosphorylation.

We have identified ABI1 as a possible AMT1 regulator in an initial screen for reduced ammonium susceptibility during hypocotyl elongation growth. The screen has led to the discovery of the knock-down line *abi1-2* (Fig. S1a/b), which shows increased tolerance to toxic methylammonium and ammonium in the hypocotyl elongation assay (Fig. S2a/b). Whereas the hydroponic growth of the *abi1* plants under standard conditions is indistinguishable from that of wild-type plants, the *abi1* plants exhibit reduced N-concentration in the shoot (Fig. S3). This suggests the possible direct effect of ABI1 on nitrogen uptake.

The increased hypocotyl elongation in the dark was reversed in complementation lines, which mimicked the wild-type phenotype (Fig. 1a). We conducted an ammonium uptake assay in order to create a link between hypocotyl growth and uptake of ammonium. This short-term uptake experiment with 0.5 and 5 mM ^15^NH_4_^+^ led to the observation of reduced ammonium uptake by *abi1* plants in high-affinity uptake conditions (0.5 mM ^15^NH_4_^+^) and a tendency of uptake reduction under low-affinity 5 mM conditions (Fig. 1c/d). Nevertheless, the differences between all lines were more pronounced at 0.5 mM ammonium compared with 5 mM ammonium. This implies an involvement of AMT1-mediated high-affinity ammonium transport in the effects of the *abi1* mutation, whereas under low-affinity conditions, apoplastic flow is most likely to overlie this effect on AMT1 activity.

Expression analysis (Fig. S5) and confocal laser scanning microscopy (Fig. S6) have excluded that lower AMT1 expression or altered subcellular localization away from the plasma membrane is responsible for the reduced ammonium uptake in the *abi1* line. Therefore, we concluded that the AMT1 phosphorylation status should be directly affected. To confirm this hypothesis, we first tested for an interaction of ABI1 with CIPK23 and with the AMT1 transporters, a prerequisite for phosphorylation. Interestingly, protein interaction studies involving the use of endogenous promotors *in planta* have revealed that ABI1 interacts with CIPK23, in addition to both AMT1;1 and AMT1;2 (Fig. 2a). We therefore hypothesize an additive effect of ABI1 on ammonium uptake via the inhibition of CIPK23 and the direct dephosphorylation of transporters (Fig. 5). As for other PP2C-As (Lee et al. 2007), ABI1 also interacts with CIPK23 and the regulated transporters in yeast, again indicating that it acts in the dephosphorylation of both proteins (Fig. 2c). In an attempt to restrict the possible ABI1-dephosphorylation sites in the AMT1 C-termini, we tested the interaction of ABI1 with deletion mutants of the AMT1s. For an AtAMT1;1 mutant in which only the conserved threonine dephosphorylation site remained, we still observed interaction between transporter and phosphatase (Fig. 2b/c), although the weak interaction was further reduced in a similar AtAMT1;2 mutant. This finding is still in agreement with the idea that ABI1 interacts and possibly dephosphorylates the conserved phosphorylation site. For AtAMT1;2, this result raises the question as to whether the C-terminus of AMT1;2 is itself dephosphorylated by the phosphatase or whether it only serves as a structure to facilitate protein interaction.

**Figure 5.**
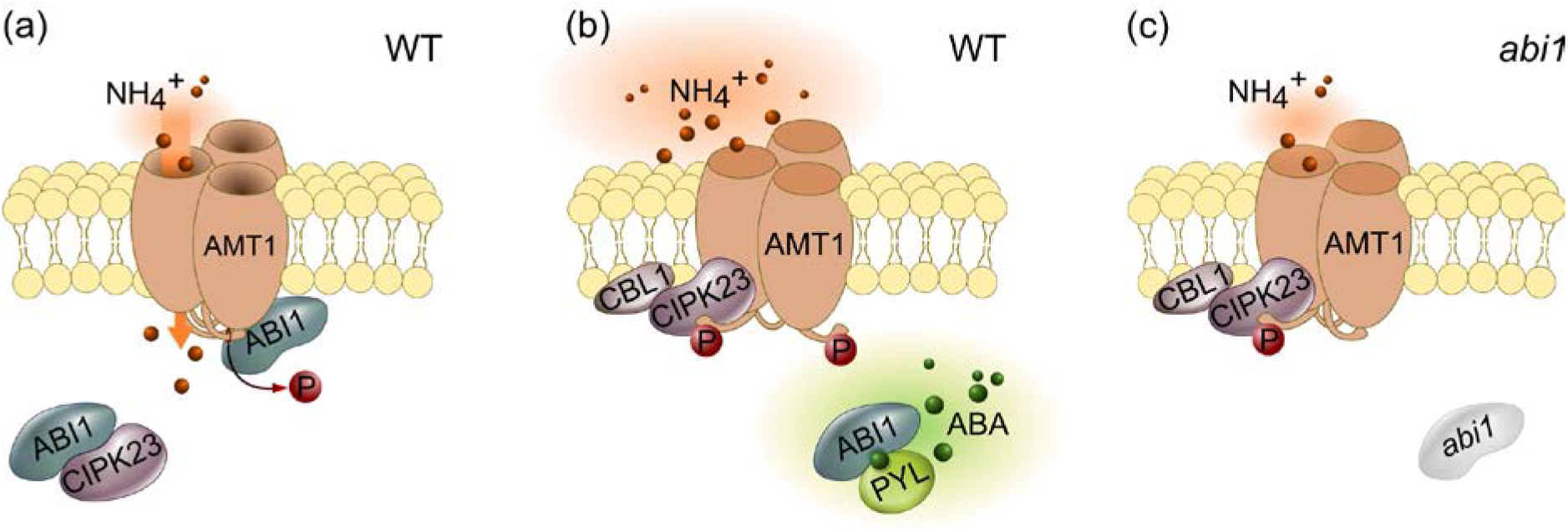
Proposed model of ammonium uptake regulation by ABI1 and CIPK23. **(a)** Under ample ammonium conditions, ABI1 will bind and inactivate CIPK23 while dephosphorylating and activating AMT1;1 and AMT1;2. **(b)** Under conditions of other types of abiotic stress and ammonium toxicity, plants produce active ABA by ABA-glycosylester deglycosylation. ABA receptors capture ABI1 and form a PYL/ABA/ABI1 complex. CIPK23 becomes released and active, phosphorylating and inactivating the AMT1 enabling the plant to avoid ammonium uptake and toxicity. **(c)** Higher AMT1 phosphorylation and reduced ammonium uptake capacity, even under low ammonium conditions in *abi1* knockdown plants.

The phosphorylation status of the AMT1s correlated well with the data from ^15^NH_4_^+^ uptake. Under nitrogen starvation, the *abi1* plants showed the highest amount of AMT1 phosphorylation (Fig. 3a) resulting in reduced ammonium uptake in comparison with all other lines (Fig. 1c). In contrast to the first time point, all lines exhibited an increase in phosphorylation at the second time point (2.5 hours after ammonium shock, Fig. 3b). As expected, this also affected ammonium uptake, which was generally reduced after ammonium shock and showed no significant differences between the lines. An additional five hours were required until the phosphorylation pattern of the four lines resembled the result following nitrogen starvation (Fig. 3d). The phosphorylation status after 5 hours showed that dephosphorylation of the AMT1s occurred faster in the “ABI1 overexpressing” complementation lines than in the wild type, whereas it was absent in the *abi1* plants (Fig. 3b and Fig. S5). Additional to demonstrating the regulation of CIPK23, these data agree with a direct AMT1 dephosphorylation function of ABI1.

We therefore conclude that, under low ammonium, non-stressed conditions, ABI1 binds to and inactivates CIPK23, whereas it can simultaneously dephosphorylate and keep AMT1s active (Fig. 5a). High external ammonium conditions stimulate a fast and strong ABA increase in the plant roots. This ABA is sensed by PYL ABA-receptors, which bind and inactivate ABI1 (Rodriguez et al. 2019). ABI1 inactivation liberates CIPK23, which in turn binds to and phosphorylates AMT1. This phosphorylation then shuts down AMT1 transport activity to reduce the ammonium toxicity effects (Fig. 5b). In the *abi1* knockdown plants, CIPK23 remains hyperactive, leading to a reduced ammonium susceptibility and higher AMT1 phosphorylation, plus reduced ammonium uptake, even under control conditions (Fig. 5c). On the one hand, plants thus sense ammonium toxicity in the same way as other abiotic stresses and use the ABA signalling cascade to integrate this stress with their ammonium nutrition. On the other hand, nutritional acclimatization as a response to abiotic stresses is probably also mediated by ABA suggesting that ABA signalling limits nutrient (ammonium) uptake under abiotic stress. This might be a general mechanism because nutrient demands are not only lower when growth is impaired by stress, but may also specifically act on ammonium transport. Investigations into whether other stresses have an effect on ammonium uptake and whether the ABA/ABI1-dependent CIPK23 regulation network also influences the transport of other ions, such as nitrate and potassium, should be of future interest.

## Acknowledgements

This work was funded by the German Research Foundation (DFG) in grants to Dr. Neuhäuser (NE 1727/2-2). We thank D. Schnell for help with the hydroponic cultures, E. Dachtler for IRMS measurements, and C. Haake for N-content measurements. We thank P. Jones for English editing.

## Author Contribution

P.G., R. P-M., T.I., N.S., U. L. and B.N. designed the experiments. P.G., R. P-M., T. I., J. M., N.M. and B. N. performed the experiments. P.G., R. P-M., T.I. and B.N. evaluated and interpreted the data. P.G, T.S. and B.N. prepared the figures. P.G. and B.N. wrote the manuscript. P.G., U.L., N.S., T.S. and B.N. revised the manuscript. B.N. obtained the funding and is the corresponding author.

## Data Availability

All data are part of the manuscript.

## Supplementary Material

**Figures S1.**
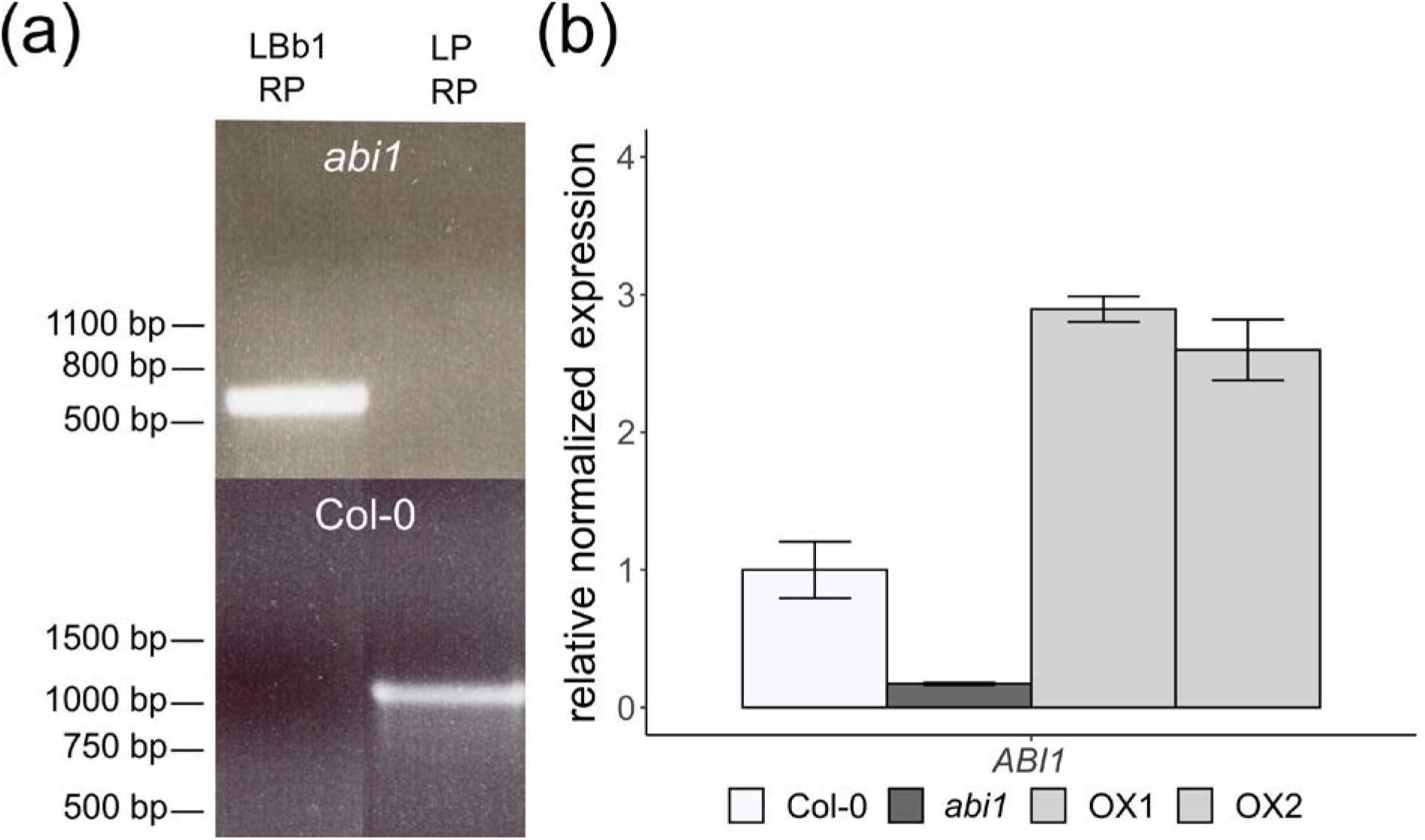
Genotyping of the *abi1-2* and complementation (OX) lines. **(a)** Genotyping of the *abi1-2* (NASC Nr.: N655606; SALK_072009C) line revealing a homozygous T-DNA insertion in *ABI1* (AT4G26080), upper part. Col-0 negative control, lower part. **(b)** Relative normalized *ABI1* expression in roots of Col-0, *abi1-2* and the complementation (endogenous promotor) line plants.

**Figure S2.**
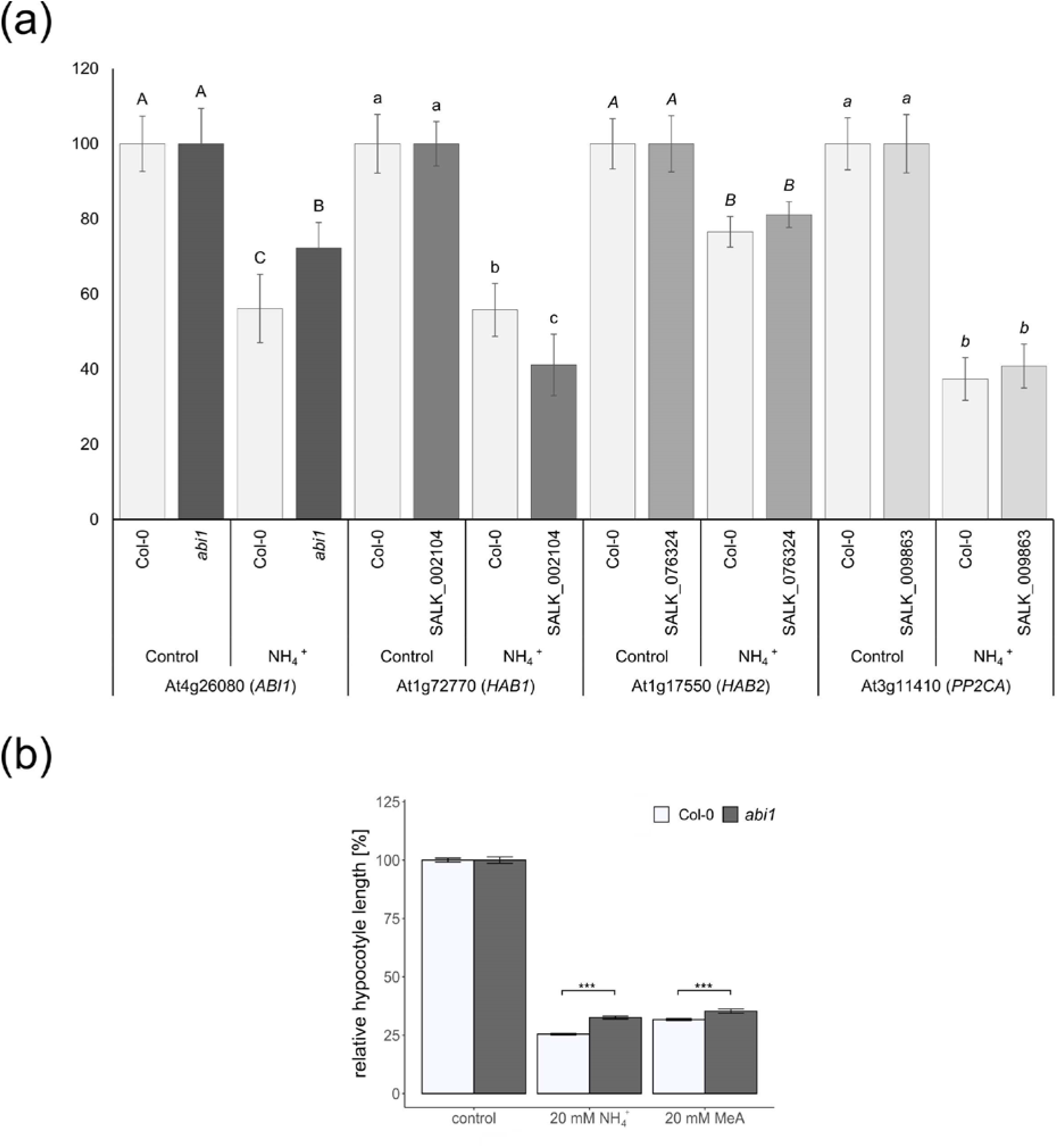
Reduced ammonium toxicity in the *abi1-2* knockdown line. **(a)** Relative normalized hypocotyl length of etiolated seedlings of PP2C class A phosphatases. Data from the original screen normalized to the control treatment. Different letters indicate statistical significance (n ≥ 20; p < 0.01). **(b)** Relative normalized hypocotyl length of etiolated Col-0 and *abi1-2* seedlings grown on control media or media containing 20 mM NH_4_^+^/methylammonium. Data are shown as means normalized to the control ±SEM (n ≥ 200; p < 0.001).

**Figure S3.**
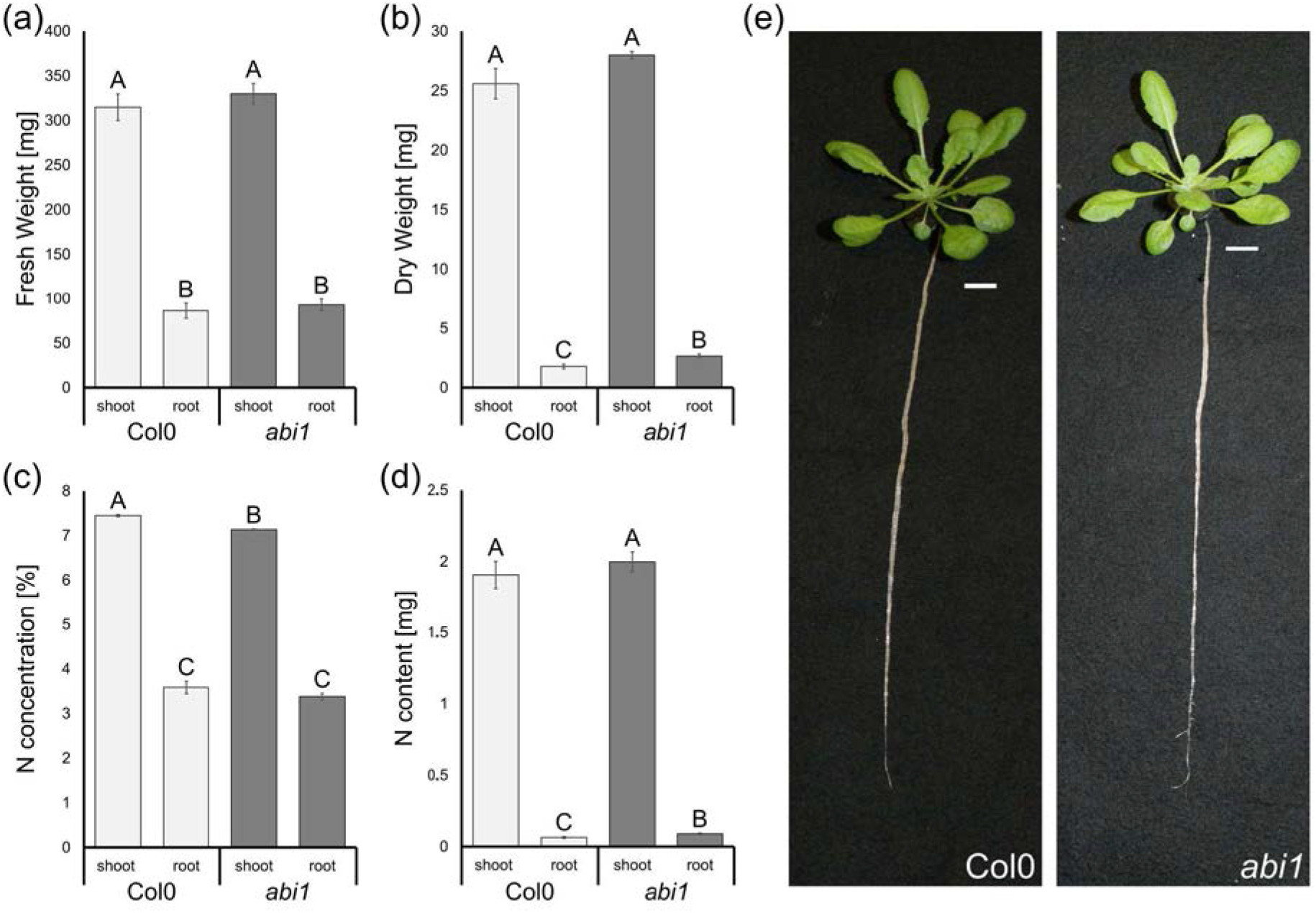
Minor growth effect of the *ABI1* knockout under non-stressed conditions but slightly reduced N-concentrations in the shoots of *abi1* plants. **(a)** Fresh weight of roots and shoots of 6-week-old Col-0 and *abi1* plants. **(b)** Dry weight of roots and shoots of 6-week-old Col-0 and *abi1* plants. **(c)** Nitrogen concentration in % (w/w) in shoots and roots of 6-week-old Col-0 and *abi1* plants. **(d)** Nitrogen content in the roots and shoots of 6-week-old Col-0 and *abi1* plants. **(e)** Images of representative 6-week-old Col-0 and *abi1* plants grown under hydroponic culture. Data are shown as means ±SEM (n = 20).

**Figure S4.**
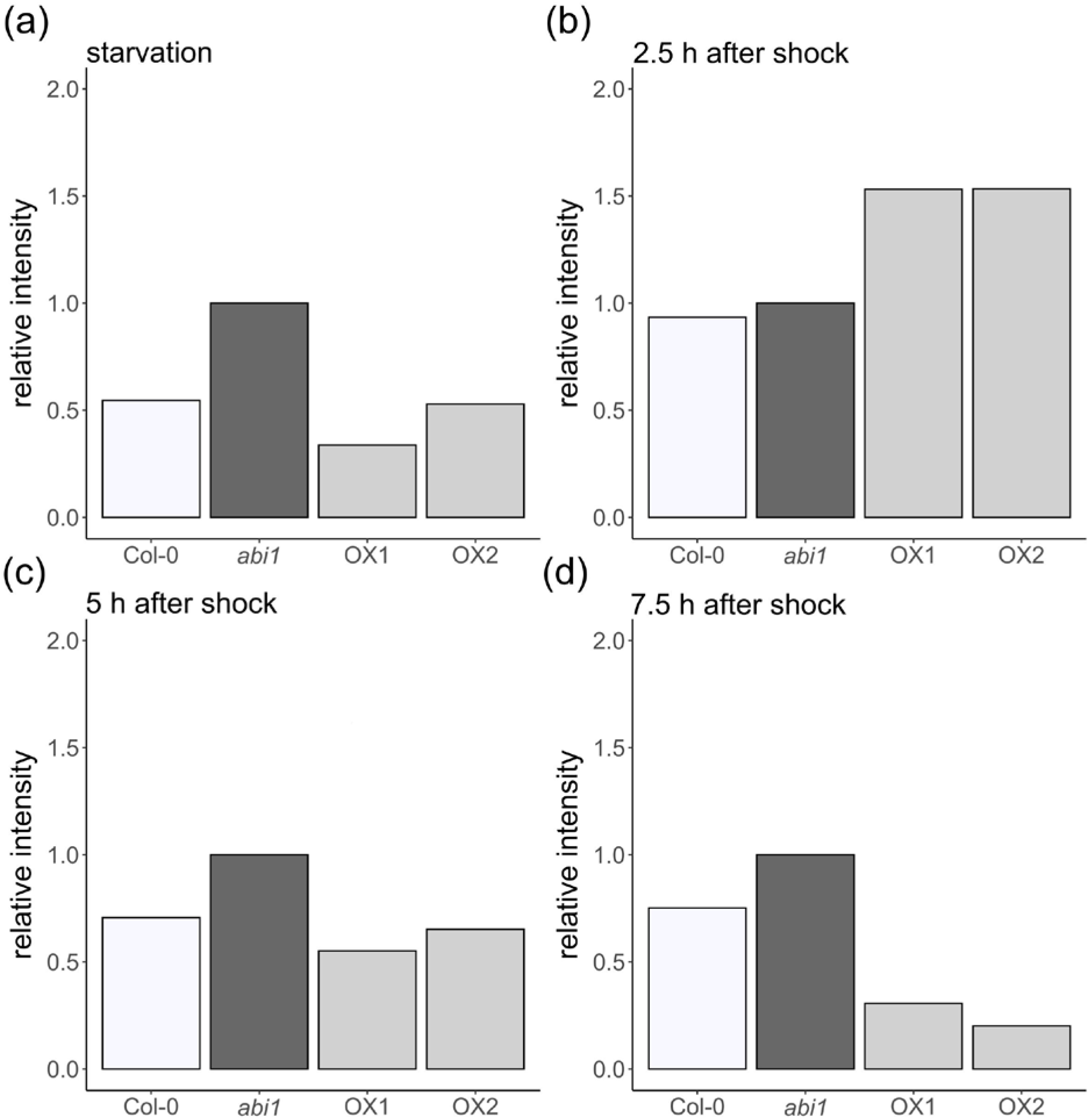
Quantification of ABI1 effects on phosphorylation status of conserved AMT1 C-termini. Protein gel blot analysis of total root protein extract from Col-0, *abi1* and complementation line plants by using a phosphorylation-specific antibody detecting the conserved AMT1 C-terminus shown in Figure 3 was quantified by ImageJ. Analysis of the two other replicates yielded similar results.

**Figure S5.**
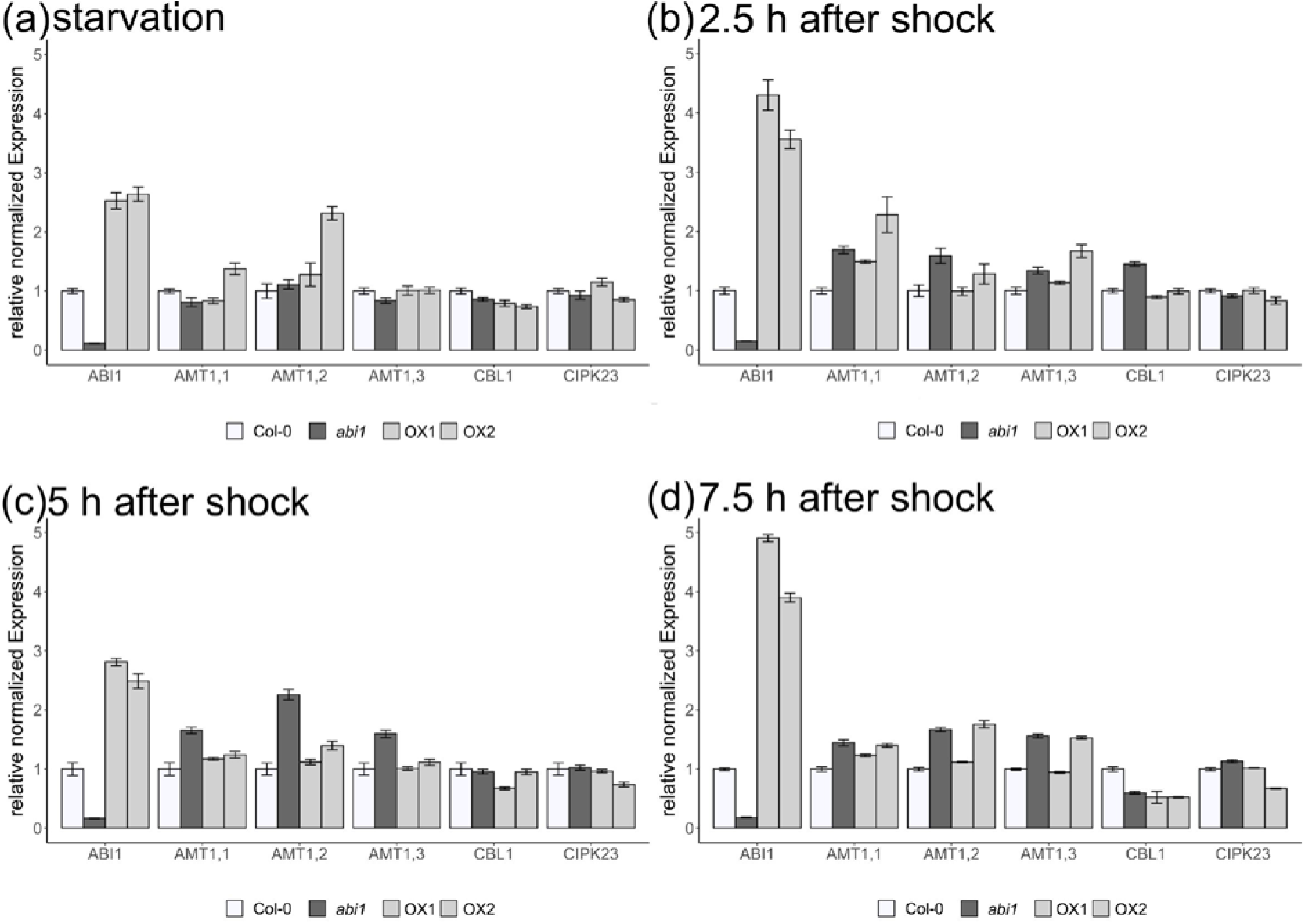
*AMT1* expression is not reduced in *abi1* mutants. Gene expression for *AMT1*s and their regulatory genes *CBL1, CIPK23* and *ABI1* in roots of 6-week-old Col-0, *abi1* and the two complementation lines. After starvation **(a)** and after 2.5 h **(b)**, 5h **(c)** and 7.5 h **(d)** of starvation after a 5-min ammonium shock.

**Figure S6.**
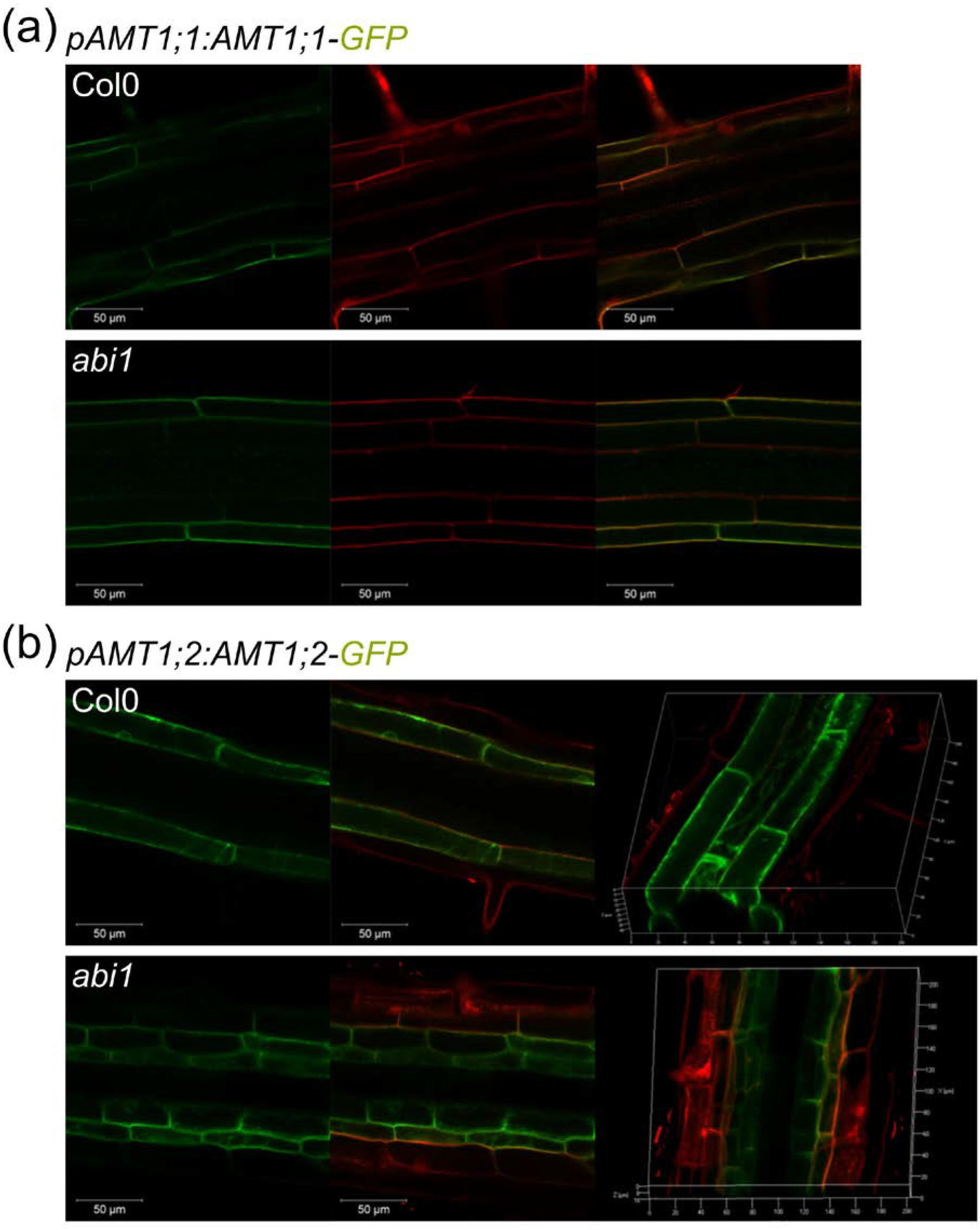
Localization of AMT1;1 and AMT1;2 is unaffected on a Col-0 or *abi1* background. Fusion construct of pAMT1::AMT1-GFP and **(a)** AMT1;1 or **(b)** AMT1;2 expressed on a wild-type or *abi1* background; fluorescence was monitored by laser scanning confocal microscopy. In (a) from left to right: GFP fluorescence, propidium iodide, overlay. In (b) from left to right: GFP fluorescence, overlay with propidium iodide, 3D view.

**Figure S7.**
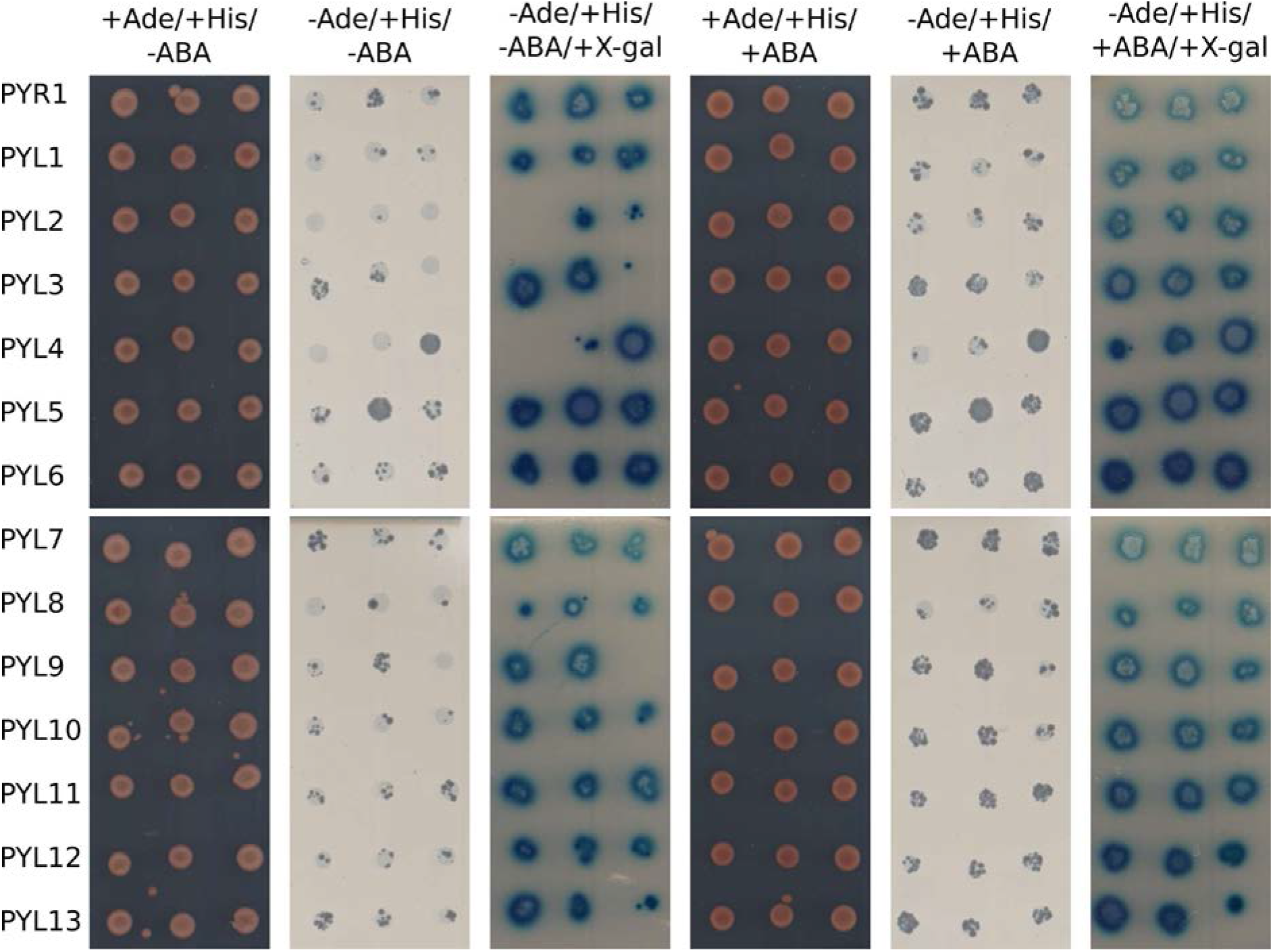
Protein interaction of ABI1 with protein family of PYR/PYL ABA receptors. ABI1 is fused to the C-terminal part of ubiquitin, whereas the interaction partners are fused to the N-terminal part of ubiquitin. Growth on media without adenine and blue staining by X-Gal overlay indicates protein interaction. From left to right, control without ABA, media without adenine and ABA, X-Gal overlay of media without adenine and ABA, control with ABA, media without adenine and with ABA, X-Gal overlay of media without adenine and with ABA.

**Table S1.** T-DNA insertion lines used in the initial screen. The number of the Arabidopsis stock centre (www.arabidopsis.info), the identification number of the T-DNA insertion line, the number of the gene associated with this insertion line, the gene family of the phosphatase and the name of the phosphatase are given.

| NASC number | SALK number | AGI Code | Gene family | Gene name |
| --- | --- | --- | --- | --- |
| 307140 | GK-065G03.02 | At1g07160 | PP2C | AtPP2C2 |
| 308984 | GK-316H10.04 | At4g27800 | PP2C | AtPP2C57/AtPPH1 |
| 315124 | GK-072G03.04 | At1g10430 | PP2A | AtPP2A1 |
| 653077 | 020722C | At4g11240 | PP1 | AtTOPP7 |
| 653085 | 021793C | At3g06270 | PP2C | AtPP2C35 |
| 654391 | 128186C | At4g16580 | PP2C | AtPP2C55 |
| 655403 | 029481C | At1g18030 | PP2C | AtPP2C8 |
| 655606 | 072009C | At4g26080 | PP2C | AtPP2C56/AtABI1 |
| 656121 | 005558C | At5g51760 | PP2C | AtPP2C75 |
| 656211 | 016641C | At3g12620 | PP2C | AtPP2C38 |
| 656223 | 017899C | At5g66720 | PP2C | AtPP2C80 |
| 656230 | 019305C | At2g20630 | PP2C | AtPP2C20/AtPPC3 |
| 656443 | 045433C | At3g05580 | PP1 | AtTOPP9 |
| 656660 | 076324C | At1g17550 | PP2C | AtPP2C7/AtHAB2 |
| 656851 | 108282C | At5g59220 | PP2C | AtPP2C78 |
| 657084 | 146020C | At4g31750 | PP2C | AtPP2C59/AtWIN2 |
| 657124 | 151946C | At5g55260 | PP4/PPX | AtPPX2 |
| 657178 | 025713C | At4g27800 | PP2C | AtPP2C57/AtPPH1 |
| 658262 | 061302C | At3g16800 | PP2C | AtPP2C41 |
| 658526 | 016135C | At3g17250 | PP2C | AtPP2C43 |
| 659308 | 013178C | At1g69960 | PP2A | AtPP2A5 |
| 659594 | 146162C | At2g34740 | PP2C | AtPP2C28 |
| 659830 | 042824C | At5g27930 | PP2C | AtPP2C73/AtPPC6 |
| 659855 | 048861C | At5g27930 | PP2C | AtPP2C73/AtPPC6 |
| 659861 | 049725C | At5g55260 | PP4/PPX-type | AtPPX2 |
| 660122 | 115494C | At5g02400 | PP2C | AtPP2C66/AtPLL2 |
| 660233 | 143218C | At1g17550 | PP2C | AtPP2C7/AtHAB2 |
| 661958 | 027747C | At3g06270 | PP2C | AtPP2C35 |
| 662121 | 036920C | At3g12620 | PP2C | AtPP2C38 |
| 662642 | 060018C | At1g07160 | PP2C | AtPP2C2 |
| 663581 | 105978C | At2g20630 | PP2C | AtPP2C20/AtPPC3 |
| 663774 | 117607C | At3g05580 | PP1 | AtTOPP9 |
| 664102 | 136570C | At1g43900 | PP2C | AtPP2C11 |
| 664699 | 076309C | At4g26080 | PP2C | AtPP2C56/AtABI1 |
| 665105 | 002104C | At1g72770 | PP2C | AtPP2C16/AtHAB1 |
| 666967 | 106575C | At2g34740 | PP2C | AtPP2C28 |
| 667683 | 003931C | At5g06750 | PP2C | AtPP2C68 |
| 667710 | 006360C | At1g78200 | PP2C | AtPP2C17 |
| 667834 | 015078C | At3g62260 | PP2C | AtPP2C49 |
| 668879 | 113717C | At4g33500 | PP2C | AtPP2C62 |
| 668957 | 122669C | At5g02400 | PP2C | AtPP2C66/AtPLL2 |
| 669012 | 127487C | At4g28400 | PP2C | AtPP2C58 |
| 669372 | 011182C | At1g69850 | PP2C | AtPP2C75 |
| 669553 | 033011C | At2g29380 | PP2C | AtPP2C24 |
| 670093 | 123000C | At3g09400 | PP2C | AtPP2C36/AtPLL3 |
| 671069 | 003944C | At1g22280 | PP2C | AtPP2C9 |
| 671261 | 014358C | At4g33500 | PP2C | AtPP2C62 |
| 672391 | 025675C | At3g62260 | PP2C | AtPP2C49 |
| 672967 | 095052C | At5g51760 | PP2C | AtPP2C75 |
| 673054 | 100800C | At1g78200 | PP2C | AtPP2C17 |
| 673072 | 102599C | At1g59830 | PP2A | AtPP2A2 |
| 673122 | 106442C | At1g09160 | PP2C | AtPP2C5 |
| 673629 | 002822C | At1g09160 | PP2C | AtPP2C5 |
| 673670 | 005240C | At2g25620 | PP2C | AtPP2C22 |
| 673778 | 009279C | At1g03590 | PP2C | AtPP2C1 |
| 673798 | 010368C | At5g53140 | PP2C | AtPP2C76 |
| 674413 | 034856C | At5g59160 | PP1 | AtTOPP2 |
| 675718 | 072696C | At1g69850 | PP2C | AtPP2C75 |
| 676774 | 123840C | At1g47380 | PP2C | AtPP2C12 |
| 676924 | 130437C | At1g07630 | PP2C | AtPP2C4/AtPLL5 |
| 678094 | 009863C | At3g11410 | PP2C | AtPP2C37/AtPP2CA |
| 678792 | 088465C | At3g17250 | PP2C | AtPP2C43 |
| 679192 | 143298C | At4g33920 | PP2C | AtPP2C63 |
| 679596 | 044162C | At1g07630 | PP2C | AtPP2C4/AtPLL5 |
| 823387 | SAIL_552_G10 | At1g22280 | PP2C | AtPP2C9 |
| 833138 | SAIL_742_A05 | At3g02750 | PP2C | AtPP2C33 |
| 840122 | SAIL_891_C01 | At3g23360 | PP2C | AtPP2C44 |
| 850628 | WiscDsLox293-296invD18 | At2g39840 | PP1 | AtTOPP4 |
| 851819 | WiscDsLox339E05 | At4g31750 | PP2C | AtPP2C59/AtWIN2 |
| 873621 | SAIL_378_F05 | At1g59830 | PP2A | AtPP2A2 |
| 878169 | SAIL_1151_F04 | At2g29400 | PP1 | AtTOPP1 |
| 903539 | WiscDsLoxHs037_11C | At1g64040 | PP1 | AtTOPP3 |
| 911811 | WiscDsLoxHs124_01C | At1g68410 | PP2C | AtPP2C15 |

**Table S2:**
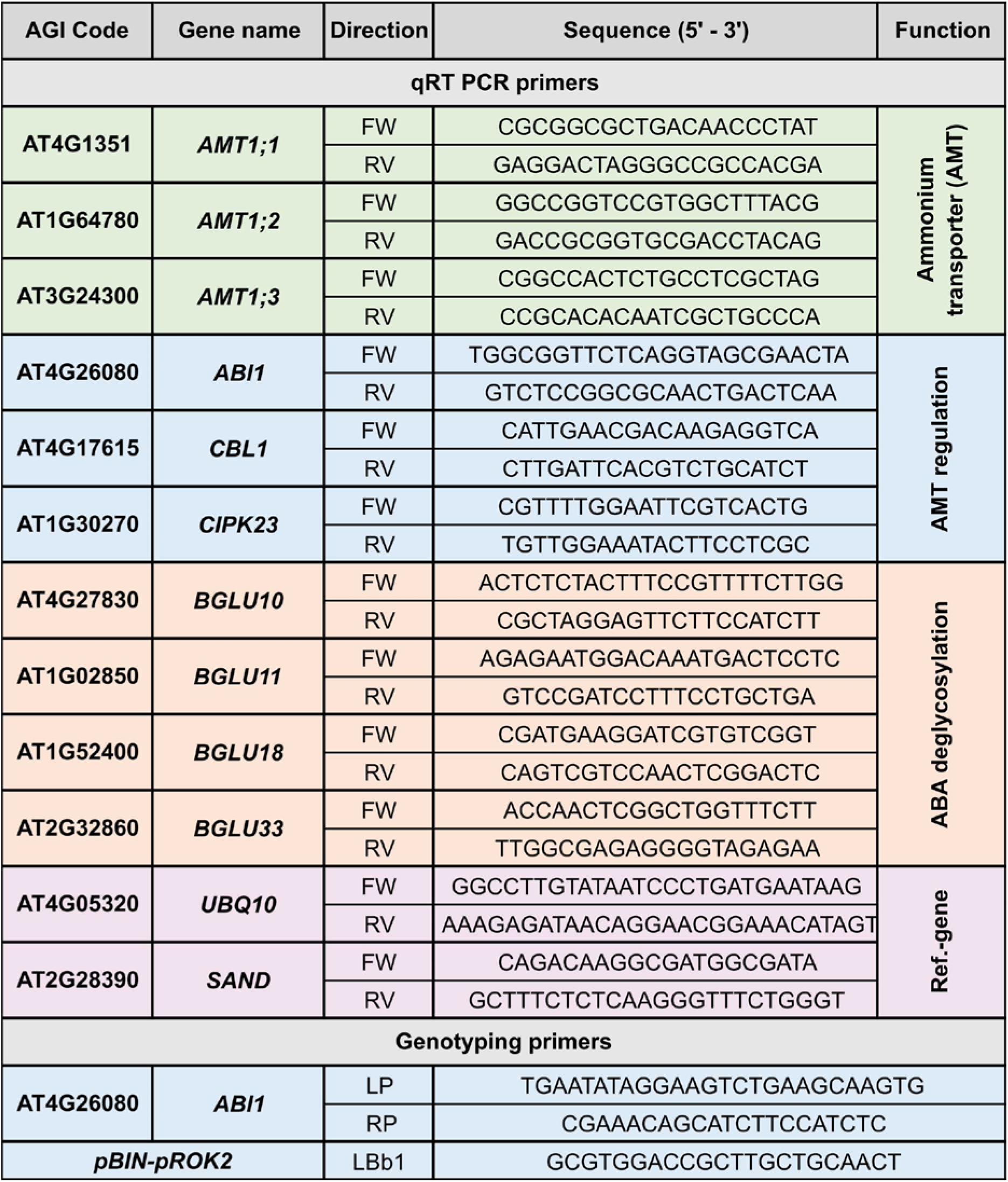
qRT-PCR and genotyping primers used in this study. The number of the gene tested in genotyping or qRT-PCR, the name of the gene, the direction of the primer corresponding to the gene, the nucleotide sequence of the primer in 5’ → 3’ direction and the function of the gene investigated in this study are given.

